# Interactions between Trypillian farmers and North Pontic forager-pastoralists in Eneolithic central Ukraine

**DOI:** 10.1101/2022.10.31.514526

**Authors:** Alexey G. Nikitin, Mykhailo Videiko, Nick Patterson, Virginie Renson, David Reich

## Abstract

The establishment of agrarian economy in Eneolithic East Europe is associated with the Pre-Cucuteni-Cucuteni-Trypillia complex (PCCTC). PCCTC farmers interacted with Eneolithic forager-pastoralist groups of the North Pontic steppe as PCCTC extended from the Carpathian foothills to the Dnipro Valley beginning in the late 5^th^ millennium BCE. While the cultural interaction between the two groups is evident through the Cucuteni C pottery style that carries steppe influence, the extent of biological interactions between Trypillian farmers and the steppe remains unclear. Here we report the analysis of artefacts from the late 5^th^ millennium Trypillian site of Kolomiytsiv Yar Tract (KYT) in central Ukraine, focusing on a bone fragment found in the Trypillian context at KYT. Diet stable isotope ratios obtained from the bone fragment place the diet of the KYT individual within the range of forager-pastoralists of the North Pontic area. Strontium isotope ratios of the KYT individual are consistent with having originated from contexts of the Sredny Stog culture sites of the Middle Dnipro Valley. Genetic analysis of the KYT individual indicates ancestry derived from a proto-Yamna population such as Sredny Stog. Overall, the KYT archaeological site presents evidence of interactions between Trypillians and Eneolithic Pontic steppe inhabitants of the Sredny Stog horizon and suggests a potential for gene flow between the two groups as early as the beginning of the 4^th^ millennium BCE.

## Introduction

Trypillian culture (5000-2750 BCE) is the eastern component of the Pre-Cucuteni-Cucuteni-Trypillia complex (PCCTC) of Eneolithic farmers of eastern Europe. PCCTC extended from the Carpathian Mountains to the Dnipro (Dnieper) River in what is now Moldova, Romania and Ukraine. PCCTC is known by over 4300 settlements and cemeteries. In the 5^th^ millennium BCE PCCTC was synchronous with such European Neolithic complexes as Lengyel, Vinca, and Kodjadermen-Gumelniţa-Karanovo VI (KGK). In the east, Trypillia, the Ukrainian branch of PCCTC, neighbored the Dnipro-Donets and Pit-Comb Ware fisher-forager groups, as well as the Sredny Stog forager-pastoralists. The Sredny Stog archaeological horizon formed in the steppe of Azov, between the Dnipro and the Don Rivers, in the 5^th^ millennium BCE. It existed, in various cultural forms, through the end of the 4^th^ millennium BCE (Kotova 2013). The Yamma cultural complex took over the steppe dominance from Sredny Stog at the end of the 4^th^ millennium BCE. Sredny Stog has been hypothesized to be the ancestral group from which the Yamna complex emerged. Sredny Stog and Yamna are viewed as an Eneolithic-Bronze Age cultural continuum that was an important source for the spread of Indo-European cultural traits across a much larger geographic region beginning in the Bronze Age (Telegin 1986).

Close contacts between Trypillia and Sredny Stog began as early as the end of the Trypillia A period (ca. 4700-4600 BCE) and continued throughout the Eneolithic (Kotova 2013). Archaeological evidence shows reciprocal influence of Sredny Stog and Trypillia on each other’s cultural development. Settlements of the Deriivka group of the Sredny Stog horizon such as Deriivka II and Molyukhiv Bugor contain evidence of agricultural practices, likely influenced by the neighboring Trypillian groups (Kotova 2013). The influence of Sredny Stog on Trypillian culture is most evident in the presence of Sredny Stog ceramic motifs in Trypillian pottery.

Archaeology and genetics point to the modern-day territory of Romania as the origin of PCCTC. Trypillian individuals from the Verteba Cave ritual site in western Ukraine share their genetic ancestry with the Eneolithic individuals of the Bodrogkeresztúr culture from Urziceni in southeast Muntenia (Mathieson *et al*. 2018). Trypillians maintained close genetic links with the farming communities of the Balkans through the end of the PCCTC existence (Immel *et al*. 2020). Archaeological studies link the origins of PCCTC with western Transylvania (Burdo 2002).

Early Trypillian sites appear in the middle Dniester area at the beginning of the 5^th^ millennium BCE, putatively reflecting eastward migration of pre-Cucuteni groups from pre-Carpathian Moldova (Burdo 2004). A plausible reason behind such a migration was climate change necessitating the exploration of new territories for arable lands. This migration has been reconstructed as having taken place in several waves. Early Trypillians, having inherited the cultural and economic adaptations of the late Neolithic cultures of the Carpathian-Danube basin, managed to significantly expand the domain of ancient agricultural societies. These early east European agrarians established the basis for the formation of what Ukrainian archaeologists refer to as the Trypillian civilization between the Dniester and the Dnipro during the second half of the 5^th^ millennium through the end of the 4^th^ millennium BCE (Videiko 2004).

After 4500 BCE Trypillia spit into several local groups, distinguished by the style of their ceramics. The emergence of Cucuteni A-B-Trypillia BI-II is likely connected with the movement of PCCTC groups out of the overpopulated Carpathian region into the forest-steppe expanses of the Dniester-Dnipro interfluve. Trypillians reached the Dnipro Valley in the second half of the 5^th^ millennium BCE. Several local Trypillian types are recognized during this period, distinguished by material culture and features of the economy. While these groups functioned autonomously, there is evidence for active interaction among them as well as with Trypillian groups to the west (Videiko & Burdo 2020). The transition to the open expanses of the Dnipro Valley also brought Trypillian groups into direct contact with steppe populations.

During the Trypillia BI-II stages, painted ceramics dominated in the western part of the expanding Trypillian domain, while the ceramics with grooved/incised décor dominated in the east. The eastern part of the Trypillian domain also adopted elements of pottery traditions from the steppe groups of the Sredny Stog cultural horizon in their kitchen pottery. Trypillian kitchenware is distinctive from other ceramic types by the composition of clay, which occasionally included crushed river shells, as well as by the shape of vessels (pots and bowls) and the method of applying ornamental compositions using stamping such as imprints of comb and pits (Figures S3, S4). This kitchenware ceramic style is known as the “Cucuteni C” type. Cucuteni C ceramics is considered an indicator of the steppe influence on Trypillian ceramics (Burdo 2016).

Around 4300 BCE, the Penezhkovska-Scherbanevska local Trypillian group was formed in the middle Dnipro Valley. The group is represented by about ten settlements, of which the two largest ones are the eponymic settlement near the village of Trypillia, over 60 hectares in size, and the Kolomiytsiv Yar Tract (KYT) near the village of Kopachiv (around 15 hectares) (Videiko & Burdo 2018) (Figure 1). Recent excavations at KYT revealed Cucuteni C ceramics in the pottery assemblage. In addition, a human long bone fragment was uncovered at the site in the Trypillian cultural layer. The presence of Cucuteni C pottery at KYT presents an opportunity to study the KYT site in the context of Sredny Stog cultural influences on Trypillia. The human osteological specimen uncovered at KYT allows the examination of diet and mobility, as well as the analysis of genetic affinities of the KYT population. In the present study, we set out to test the hypothesis of the existence of genetic interactions alongside cultural exchanges between Trypillia and Sredny Stog at Trypillian sites containing Cucuteni C pottery. Finds of human remains at Trypillian sites are exceedingly rare (Nikitin *et al*. 2010, 2017b). The osteological specimen from KYT provides exceptional opportunity to expand our understanding of subsistence patterns, mobility, and genetic landscape of the area to the south of the Carpathian Mountains during the Eneolithic.

**Figure 1.**
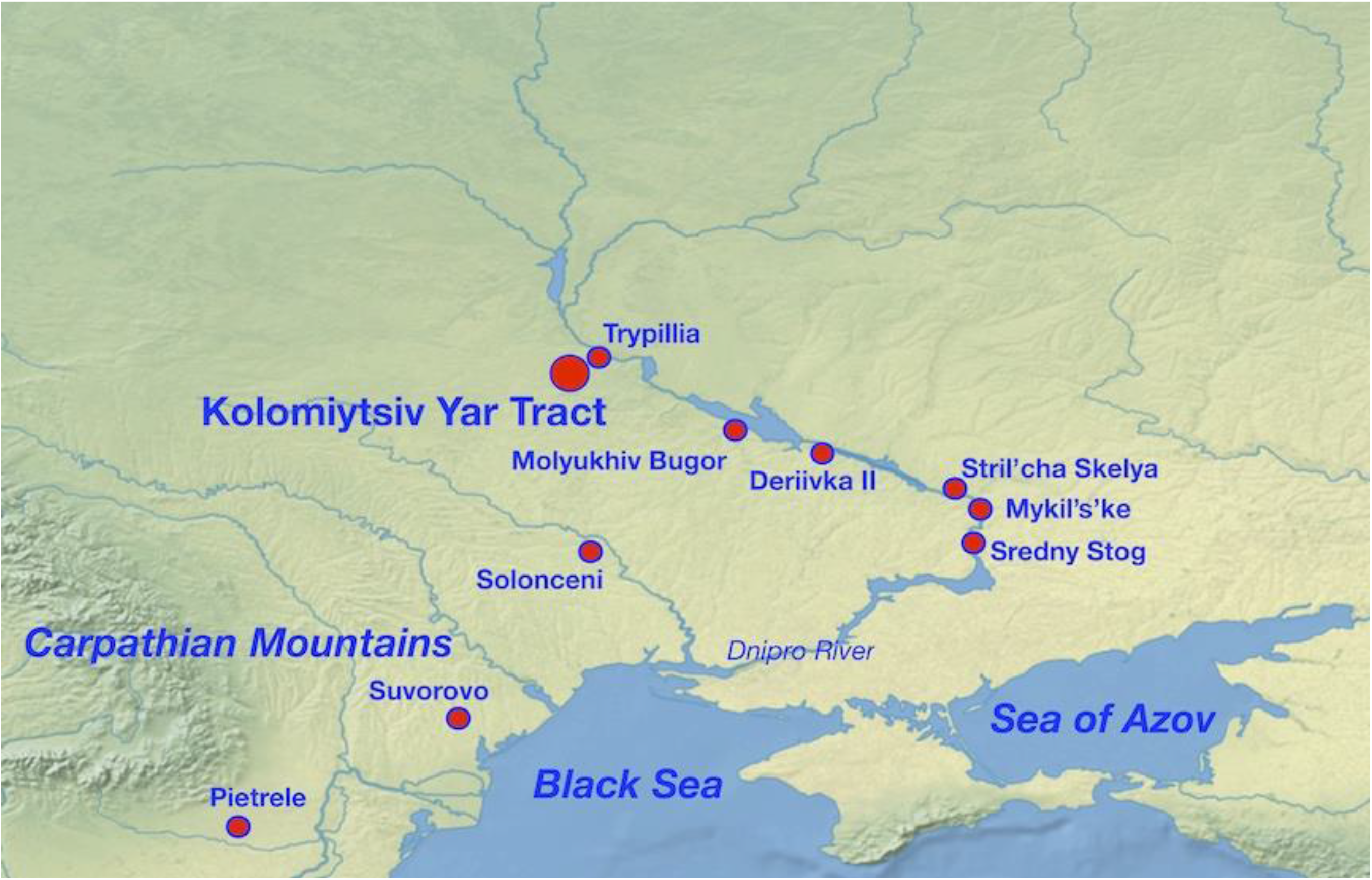
Location of the Kolomiytsiv Yar Tract, as well as Trypillia, Derrivka II, Molyukhiv Bugor, Sredny Stog, Stril’cha Skelya, Pietrele, Suvorovo, Solonceni and Mykil’s’ke archaeological sites mentioned in the text. Image from https://www.naturalearthdata.com/ (public domain),modified.

## Materials and Methods

### Kolomiytsiv Yar Tract site description

Within the modern limits of the Kyiv Region, more than two dozen settlements of stage ВІ-II are now known. Trypillian settlements of this period, in comparison with those of the later stages BII, CI and CII, are among the least studied in central Ukraine. The settlement at the Kolomiytsiv Yar Tract (KYT) is one of the easternmost among the Trypillian sites in the northern part of this region.

Kolomiytsiv Yar Tract (50.10799, 30.52610) is situated in a valley, southeast of the village of Kopachiv, Obukhiv District, Kyiv Region (Figure 1). A stream, which is the west tributary of the Stugna River, flows southwest-northeast along the bottom of the valley (Figure S5). The left, sloping bank of the stream is occupied by farm fields. The steep right bank of the stream is partially overgrown with forest. The floodplain of the stream is marshy, has a width of 100 to 50-30 m, and is partially overgrown with reeds (Figures 2, S2).

**Figure 2.**
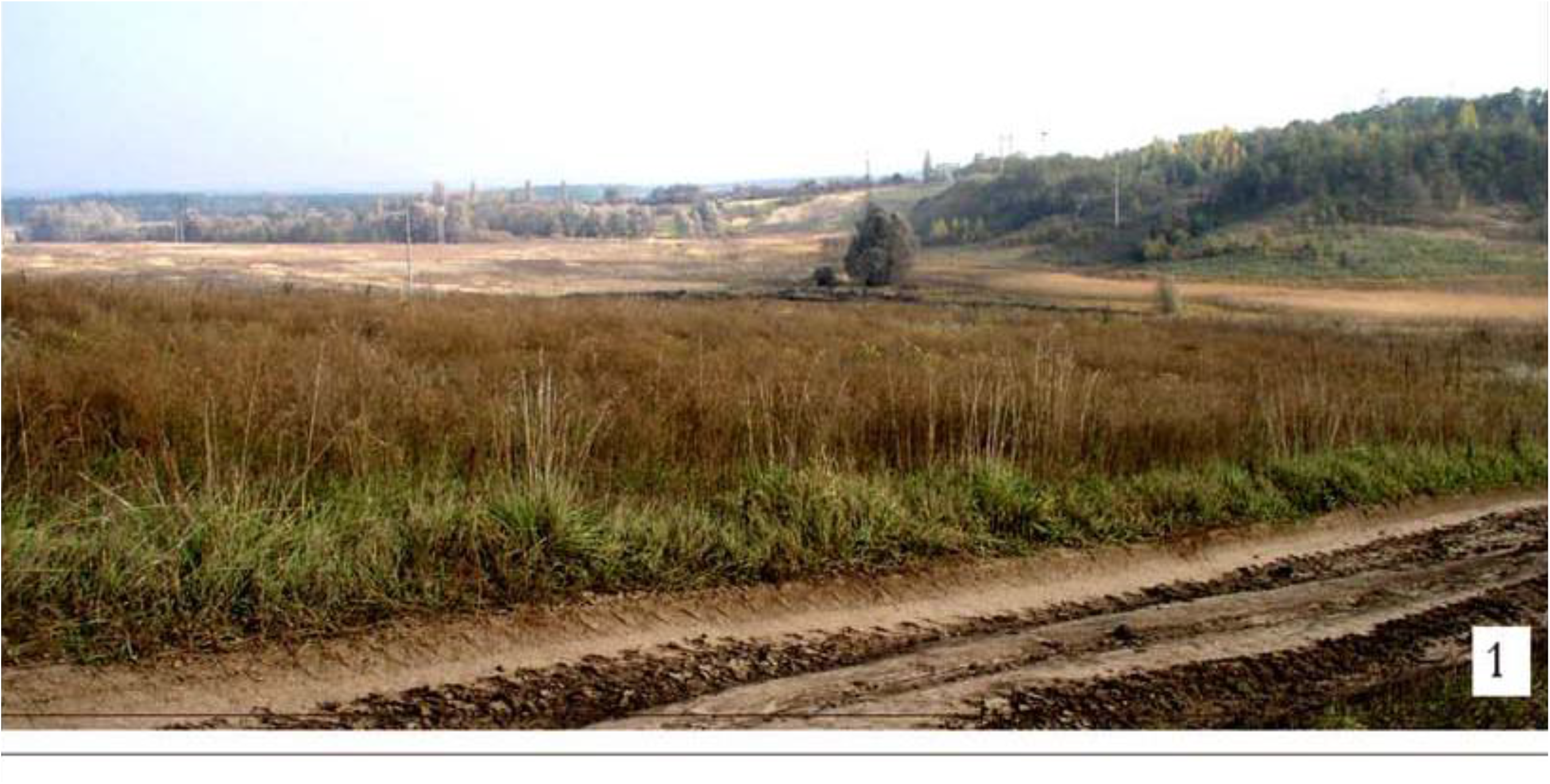
The Kolomiytsiv Yar Tract setting. Photo by M. Y. Videiko, 2006.

The KYT archaeological site was discovered in 2005 (Kvitnitskiy 2006). It was further excavated by a joint expedition of the Kyiv Regional Archaeological Museum and the Institute of Archeology of the National Academy of Sciences of Ukraine in 2006 (Videiko *et al*. 2007), 2007 (led by Maxim Kvitnicky), and 2011 (Videiko 2012). These were rescue excavations since part of the site’s cultural layer was being actively destroyed by the extraction of topsoil (chernozem) in the floodplain of a stream along which the site’s Trypillian structures were found. The site is multi-layered and includes materials from the Trypillian culture (stage BI-II), the early Iron Age, and the Medieval periods (XII-XIII centuries). Archaeological monuments, later in time than the Trypillian culture, were located in the lower part of the valley, in a 70-100 m strip along the right bank of the stream.

In cooperation with the Christian-Albrecht University, the “Human Development in Landscapes” School (Kiel, Germany), a magnetic survey was carried out on a part of the KYT’s Trypillian settlement in 2016. The survey covered about three hectares of an area of the slope of the promontory in the lower part of the valley (Videiko *et al*. 2017). As a result, underground anomalies corresponding to the remains of 19 incinerated buildings of the Trypillian culture and traces of six pits of various sizes were found (*ibid*., Fig. 1). The survey determined that the anomalies were of varying preservation - from good to severely destroyed by plowing and erosion, especially those located in areas with a considerable slope of the terrain. Field excavations confirmed the results of the magnetic survey both regarding the presence and preservation of the underground features.

Underground features representing remains of buildings were located in three rows on the sloped terrain along the stream flowing along the bottom of the valley. The top row contained the remains of 11 buildings. The second row contained the remains of five structures, and the bottom row had three. Probably most of the remains of the buildings on this site were located outside the field, where some building remains were surveyed in 2005-2007. Pit anomalies were identified near the remains of buildings. The largest pit was located opposite the end of a building under excavation 14, which also stood out for its size. The pit’s location suggests that originally these pits were used for clay extraction during dwelling construction by Trypillians.

In the heavily disturbed area of the topsoil extraction, a human long bone fragment (possibly a humerus shaft, Figure S1) was discovered in 2007. The bone fragment was found in the proximity of a burnt house (“*ploschadka*”) of Trypillian culture, explored during excavations in 2007.

### Radiocarbon dating and diet stable isotope analysis

Radiocarbon dating using Accelerated Mass Spectrometry (AMS) as well as carbon and nitrogen (δ^13^C and δ^15^N) stable isotope analyses were carried out at BETA Analytic, Miami, FL. Stable isotope measurements were obtained using a modified version of the Longin collagen extraction method (Longin 1971). Radiocarbon calibration was carried out using OxCal version 4.4 and the IntCal 20 calibration curve (Bronk Ramsey 2009; Reimer *et al*. 2020). Stable isotope analysis of carbon and nitrogen is used in archaeological investigations to examine dietary patterns. Carbon isotope ratio, δ^13^C, is a measure of the ratio of the stable isotopes of carbon, ^13^C and ^12^C. Carbon isotope ratios are measured on bone collagen and dentin, which makes it possible to determine the origin of dietary proteins from marine, terrestrial and freshwater resources. Within the terrestrial sources, δ^13^C makes it possible to differentiate between the C3 and C4 plant-based diets (Schwarcz & Schoeninger 1991). Nitrogen isotope ratio, δ^15^N, is the ratio of stable isotopes of nitrogen, ^15^N and ^14^N. The evaluation of δ^15^N makes it possible to determine the trophic level of the studied organism within a food web. Populations of different subsistence strategies can be distinguished by their stable isotope signature (Schwarcz & Schoeninger 1991).

### Strontium isotope analysis

Strontium isotope ratio analysis of the KYT bone and soil specimens was carried out at the University of Missouri Research Reactor (MURR). The soil sample was powdered in an agate mortar and calcined at 550°C for four hours. An aliquot of about 0.1 g was transferred to a PFA vial and dissolved in 24N HF - 14N HNO_3_ (4ml and 1ml, respectively) on a hot plate at 125°C for 48hrs. The solution was then evaporated and the residue re-dissolved in 6ml of 6N HCl at 125°C for 48hrs. The solution was evaporated at 90°C, and the residue was re-dissolved in 2ml of ^14^N HNO_3_ at 125°C and evaporated.

The bone sample was mechanically cleaned using a microdrill equipped with a bristle brush. It was then chemically cleaned using an ultra-sonic bath and a succession of 0.1N acetic acid for 30 minutes, mQ water for 15 minutes, and 0.1N acetic acid for 15 minutes. The sample was thoroughly rinsed with mQ water after each cycle. The bone was then leached for seven hours in 5% acetic acid, rinsed with mQ water, and dried at 105°C in the drying oven. A fragment of about 40-50 mg was dissolved in a PFA vial using 7N HNO_3_ on a hot plate at 110°C for 24hrs. The solution was then evaporated at 90°C.

The dry residues of the bone and the soil samples were re-digested in 2ml 7N HNO3 and the Sr was extracted using a protocol adapted from (de Muynck *et al*. 2009). The Sr solutions were evaporated, and the residue dissolved in 0.05N HNO_3_ before analysis.

The Sr isotopic analysis was conducted on a Nu Plasma II (Nu Instruments) multi-collector – inductively coupled plasma – mass spectrometer in operation at MURR. Both the samples and the SRM987 Sr isotopic standard solutions were prepared to obtain a Sr concentration of about 150 ppb. The measured ratios were corrected for the isobaric interference of ^87^Rb, ^86^Kr, and ^84^Kr, and for mass bias using the iterative approach and a value of 0.1194 for ^86^Sr/^88^Sr natural ratio. The SRM987 was measured multiple times (n=11) and the value obtained for the ^87^Sr/^86^Sr were 0.71022 ± 0.00004 (2sd). The values obtained for the samples were corrected by standard bracketing using the accepted value of 0.710248 for the ^87^Sr/^86^Sr (Thirlwall 1991).

### Genetic analysis

We carried out the first steps of ancient DNA analysis in a dedicated clean room at Harvard Medical School, following previously established protocols to minimize contamination including working in positively pressured rooms decontaminated with ultraviolet light, and use of protective clothing by the technicians handling the remains. We took a sample of xx mg of cortical bone powder using a dental drill. We extracted DNA using a methodology optimized to retain short and degraded molecules (Dabney *et al*. 2013; Korlević *et al*. 2015; Rohland *et al*. 2018), and converted the extracted DNA into a barcoded, double-stranded library (Rohland & Reich 2012). We enriched the library in solution for sequences overlapping at least 1.2 million single nucleotide polymorphisms (Fu *et al*. 2015; Rohland *et al*. 2022). We sequenced on Illumina instruments using 2×76bp sequences. We processed the data bioinformatically and mapped to the human reference genome hg19 as described in previous studies, removed duplicated molecules based on matching barcodes and start and stop position in the mapped genome (e.g. (Mathieson *et al*. 2015). We represented each targeted position by a single randomly chosen sequence, which produce a total of 357,145 targeted SNPs on chromosomes covered by at least one sequence. The sequence data gave no evidence of contamination based on polymorphism in its mitochondrial DNA using contamMix (Fu *et al*. 2013) (the 95% confidence interval for matching to the consensus haplogroup of U4b1b2 was 98.9-100%) and had ratio of Y to X chromosome sequences consistent with female molecular sex.

## Results and Discussion

### Ceramic assemblage at Kolomiytsiv Yar Tract

Detailed analysis of the KYT ceramic assemblage is presented in (Videiko *et al*. 2017). Overall, Trypillian ceramics at the KYT site belongs to the stage ВІ-II of Trypillia culture period. The ceramic assemblage uncovered at the site is typical for this stage in the Middle Dnipro region and is represented by the so-called “kitchenware” and “tableware” types. A characteristic feature of the ceramics from KYT is the addition of crushed shells to the clay of some vessels with a carved ornament, as well as the processing of their inner surface with a tool that leaves stripes - combs, which is typical for ceramics of the Cucuteni C type (Figures S3, S4). A detailed comparative analysis of ceramics of the Cucuteni C type from the settlements of Trypillia and Sredny Stog convincingly shows that Trypillian settlements contain imported (Sredny Stog) and locally made ceramics with shell admixture (Tsvek & Rassamakin 2002). Finds of Cucuteni C ceramics at KYT are of local production. Cucuteni C pottery at KYT parallels the Sredny Stog ceramics of Sredny Stog II, Stril’cha Skelya and Molyukhiv Bugor (Figure S4). The ceramic assemblage at KYT containing Cucuteni C ceramics made on site suggests the presence of carriers the Sredny Stog pottery making techniques at KYT.

### Absolute dating and dietary isotopes of the KYT individual

Kolomiytsiv Yar Tract human osteological specimen was directly dated to 5170±30 BP (Table 1). After calibration, this corresponds to 4049-3820 calBCE (95.4% probability). Thus, radiocarbon analysis confirms the placement of the specimen within the Trypillian BI-BII period.

**Table 1.**
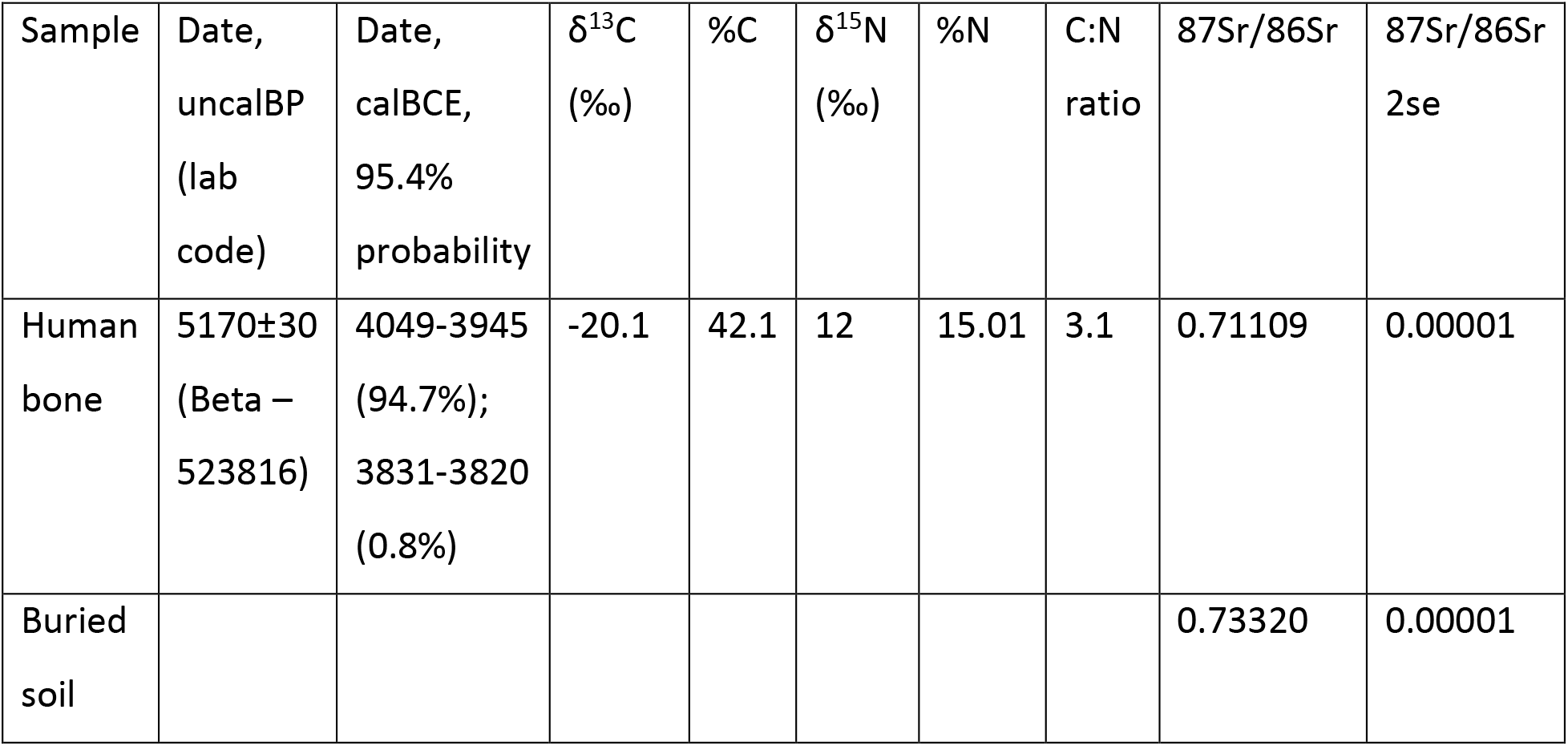
Isotope data of the Kolomiytsiv Yar Tract samples.

The δ^15^N of 12.0‰ suggests the potential presence of Freshwater Reservoir Effect (FRE) from aquatic dietary sources (Lillie *et al*. 2009, 2011). The δ^15^N terrestrial baseline is between 9.5 and 11‰. Past 11.5‰, the FRE can influence the date up to 200 years (M.C. Lillie, personal communication, 2021). Considering the δ^15^N of the KYT specimen being just over the terrestrial baseline threshold, the FRE influence on the date is likely at the lower end of the FRE influence scale. It is not possible to quantify the FRE more precisely in the absence of contemporaneous faunal remains.

Stable isotope values of the KYT specimen were plotted against the corresponding values from Ukrainian Neolithic, Eneolithic and Early Bronze Age populations for which stable isotope data are currently available (Figure 3, Table S1). Kolomyitsiv Yar values for both δ^13^C and δ^15^N were outside of the range of variation of the Trypillian farming groups from Verteba Cave (Lillie *et al*. 2018) and Prydnistryanske of the Yampil complex (Goslar *et al*. 2017).

**Figure 3.**
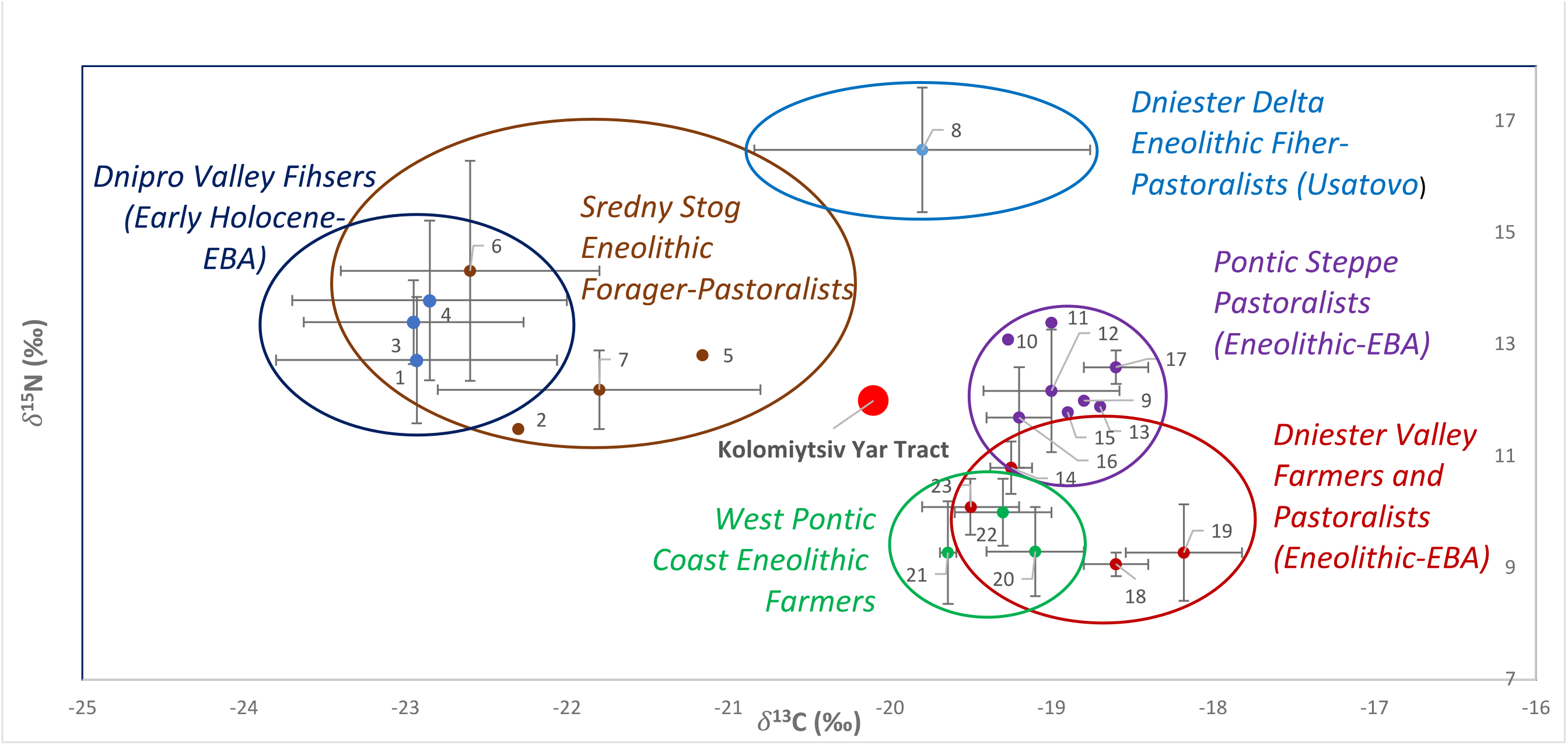
Distribution of δ^13^C and δ^15^N stable isotope ratios of Early Holocene-Bronze Age populations of the North and West Pontic area. Stable isotope ratios and corresponding publication sources are listed in Table S1. 1, Deriivka I, III; 2, Deriivka II; 3, Mykil’s’ke; 4, Yasynuvatka; 5, Oleksandria; 6, Igren 8; 7, Molyukhiv Bugor; 8, 12, Mayaki; 9, 13, Pischanka; 10, Kamʺyanka-Dniprovsʹka; 11, 17, Vinogradne; 14, Pidlisivka/Porohy; 15, Shakhta Stepna; 16, Sugokleya; 18, 19, Prydnistryanske; 20, Durankukak; 21, Smyadovo; 22, Varna; 23, Verteba Cave.

Carbon and nitrogen stable isotope ratios of the KYT specimen were comparable with diet isotope ratios of the North Pontic steppe groups. Nitrogen δ^15^N ratios of KYT were within the range of variation of the Eneolithic-EBA Pontic steppe pastoralists, Eneolithic forager pastoralists from the Sredny Stog horizon, as well as the Dnipro Valley fishers, at the lower range of δ^15^N ratio variation for the latter group (Figure 3). Carbon δ^13^C ratios of the KYT specimen were within the range of variation of the Dniester Delta Eneolithic population from the Usatovo site at Mayaki, with likely subsistence based on fishing and pastoralism, and outside the range of carbon ratio variation for steppe pastoralists. Overall, the KYT diet appears comparable to the diet of fisher-forager-pastoralists of the North Pontic steppe, but not to the diet of farmers of Eneolithic Pontic coast and adjacent forest steppe areas (Figure 3). Based on diet isotope values, we conclude that KYT comes from a population with the subsistence strategy based on foraging and/or pastoralism.

At the same time, we cannot exclude that the particular isotope ratios found in the KYT individual could be a result of some special diet. The diet of the KYT individual could also be reflecting a mixture of farming, pastoralist and foraging subsistence practices, if KYT were a forager living in a farming environment or vice versa.

The presence of Cucuteni C ceramic style at KYT suggests the interaction of the Trypillian population of KYT with and influence by Sredny Stog. The two most proximal to KYT Sredny Stog settlements on the right bank of the Dnipro River are Deriivka II and Molyukhiv Bugor (Figure 1). The Sredny Stog layer at the Molyukhiv Bugor site is dated to 3951-3640 BCE (Lillie *et al*. 2011). Radiocarbon dates from the Deriivka II settlement place it in 4436-3988 BCE (Lillie *et al*. 2009). It has been suggested that the formation of culture groups of the Sredny Stog II period (according to D. Telegin’s classification (Telegin 1973)), represented by settlements at Molyukhiv Bugor and Deriivka II, was influenced by Trypillia (Telegin 1986; Rassamakin 1999; Kotova 2013). Archaeological evidence suggests that the Sredny Stog populations at Deriivka II and Molyukhiv Bugor practiced agriculture, but their main form of subsistence was hunting and pastoralism, supplemented by freshwater fish and mollusks (Telegin 1973, 1986; Lillie *et al*. 2009; Kotova 2013; Mileto *et al*. 2017). The presence of Sredny Stog groups in the forest steppe zone indicates the movement of Sredny Stog northward during the Srednij Stog II period.

### Strontium isotope analysis

The ^87^Sr/^86^Sr ratio value measured in the KYT bone specimen was 0.71109 and the one measured in the buried soil was 0.73320 (Table 2). No samples were collected to define a strontium isotopic baseline for the purpose of the present study. The strontium isotopic composition of the bone and soil samples were compared to data from (Ventresca Miller *et al*. 2021) for estimated values of the ^87^Sr/^86^Sr ratio values across the different geological ages in Ukraine. These estimations were derived from values presented in (Voerkelius *et al*. 2010). The geological setting of the KYT site and the adjacent stream is mainly composed of sedimentary rocks and sediments from the Eocene, Oligocene and Miocene, with estimated range of ^87^Sr/^86^Sr ratio values between 0.709 and 0.711 (Ventresca Miller *et al*. 2021). Rocks of Precambrian age with estimated range of values ^87^Sr/^86^Sr ratio values between 0.712 and 0.780 (Ventresca Miller *et al*. 2021) can be found West and South within a range of 35-45 km from the KYT site. The lithologies around the KYT site are presented in detail in Figure S5.

**Table 2.** Genomic data of the Kolomiytsiv Yar Tract sample. Full DNA library report is presented in Table S2.

| Specimen/DNA lab code | Molecular sex | MtDNA haplotype | SNP hits on autosomal targets | Coverage on autosomal targets |
| --- | --- | --- | --- | --- |
| Human long bone /I7585 | Female | U4b1b2 | 76612 | 0.068193 |

The ^87^Sr/^86^Sr ratio value of the soil sample (0.73320) is higher than the range proposed for the Cenozoic substrate present at the KYT site. No strontium isotopic data on the specific lithologies developed at the site is available, which makes it difficult to assess whether the soil is representative of the substrate or if strontium from another source has contaminated the soil. The site itself, and the area drained by the adjacent stream, are not currently farmed, but agriculture is common in the region and the application of fertilizers is likely. Numerous studies have demonstrated the impact of modern fertilizers on the strontium isotopic signature of soils (e.g., (Antich *et al*. 2000; Böhlke & Horan 2000; Lottermoser 2009) and archaeological remains buried in these soils (e.g., (Bentley 2006). Available data for the strontium isotopic composition of fertilizers exhibit lower values than that of the soil sample from KYT (e.g., (Antich *et al*. 2000; Böhlke & Horan 2000; Vitòria *et al*. 2004). However, some data for K-fertilizers also demonstrates that they can present much higher values (e.g., (Borg & Banner 1996; Böhlke & Horan 2000). Additional data, including the identification of the types of fertilizers used in the region, would be required to determine if the Sr isotopic signature from the KYT soil has been affected by the use of K-fertilizers. No older rock of Proterozoic or Archean age with higher strontium ratio value is present along the stream adjacent to the site (Figure S4), and all lithologies around the site are of Cenozoic age and exhibit lower Sr ratio values than the soil sample. Further sampling around the site would be needed to define the strontium isotopic signature of the site and its immediate surroundings.

The KYT bone sample ^87^Sr/^86^Sr ratio value is slightly higher than the range of estimated values for the Cenozoic substrates. The signature in the bone could be the result of a mixture of Cenozoic substrates with an input from Precambrian rocks that present higher strontium ratio values. This would be consistent with the diet of a forager-pastoralist who would get their food from the area of Precambrian rocks to the South and West from the KYT site. However, these substrates, and regions where they can be found in association, have a broad distribution, which prevents precise association of the KYT bone specimen with a specific area or site.

The bone sample’s strontium ratio was also compared to the signature obtained from human enamel from individuals of the Eneolithic and Yamna populations of Ukraine (Gerling 2015; Ventresca Miller *et al*. 2021). The KYT individual’s strontium ratio falls within the range of values observed for the site of Sugokleya (Kirovograd Region), which is associated with the Yamna culture (Gerling 2015). More specifically, the KYT bone’s strontium ratio is close in value to that from the enamel of an individual found at the Sugokleya Kurgan Grave 10 (UK 44/45, (Gerling 2015)). The Sugokleya site is located in the vicinity of Deriivka II settlement of the Sredny Stog culture, making that area a plausible place of origin of the KYT individual. The present data, however, do not allow to unequivocally identify the place of origin of the KYT individual nor to firmly establish whether the individual was non-local to the KYT site.

The fact that bone is a porous tissue and its biogenic strontium signature is commonly affected by diagenesis/chemical exchanges within the burial environment has consequences for this research. Because the soil in immediate contact with the bone is higher than that of the bone, it is possible that the value of the bone was originally lower than the value that was measured.

Taken together, diet isotope analysis of the KYT individual suggests the forager-pastoralist subsistence type such as that of forest-steppe Sredny Stog communities of the middle Dnipro Valley. Strontium isotopic signature of the KYT individual is compatible with a large geographic area, encompassing most of the Middle Dnipro Valley, and close to the strontium isotope ratios in the vicinity of the Deriivka II settlement of Sredny Stog (Figure 1). The presence of Cucuteni C pottery at the KYT site pointing at the presence of Sredny Stog pottery makers at KYT (Videiko & Burdo 2018) and the similarity of ceramic assemblages between KYT and Sredny Stog, further support the origin of the KYT individual from a Sredny Stog population.

### Genetic analysis of the KYT specimen

To examine the genetic ancestry of the KYT individual, we extracted and examined the KYT individual’s DNA. Genetic analysis revealed that the individual was a female, carrying mtDNA haplogroup U4b1b2 (Table 2). Carriers of the U4b subclade were identified in Mesolithic and Neolithic fishers and foragers from Ukraine, the Iron Gates area of the Danube, and the Baltic coast (Mathieson *et al*. 2018). The U4 mtDNA clade was present in the Middle Dnipro Valley from the beginning of the Holocene and persisted in the North Pontic steppe through at least the late Eneolithic (Nikitin *et al*. 2017a; Mathieson *et al*. 2018; Allentoft *et al*. 2022; Mattila *et al*. 2022). Carriers of the U4b mtDNA lineage have not been identified in Trypillian remains studied to date. At the same time, a late Trypillian individual dated to 3482-3297 calBCE from the Gordineşti site in Moldova and identified as having a substantial “steppe” genetic admixture carried a U4-derived mitochondrial lineage U4a1 (Immel *et al*. 2020). On the whole-genome Principal Component Analysis (PCA), the Gordineşti individual is pulled away from the cluster of Trypillian specimens from Verteba cave, dated to ca. 3900-3600 calBCE, towards the EBA Yamna individuals from Ukraine and the Volga River region.

Whole genome analysis of the KYT specimen produced 76550 Singe Nucleotide Polymorphism (SNP) “hits” covering 6.8% of total autosomal targets (Tables 2, S2). We produced a PCA projection of ancient and modern west Eurasian genomes that included the KYT specimen (Figure 4). Many ancient samples fall outside the genetic range of modern Eurasia. The KYT sample (marked as SSX in Figure 3) falls close to Yamna pastoralists of the EBA. KYT-SSX has more ancestry from the ancient steppe than any modern West Eurasian sample, but is very distinct from Mesolithic-Neolithic hunter-gatherers, either Eastern Hunter Gatherers (EHG) or even more so samples from the Mesolithic in Serbia (Iron Gates) which are genetically primarily of Western Hunter Gatherer (WHG)-associated ancestry.

**Figure 4.**
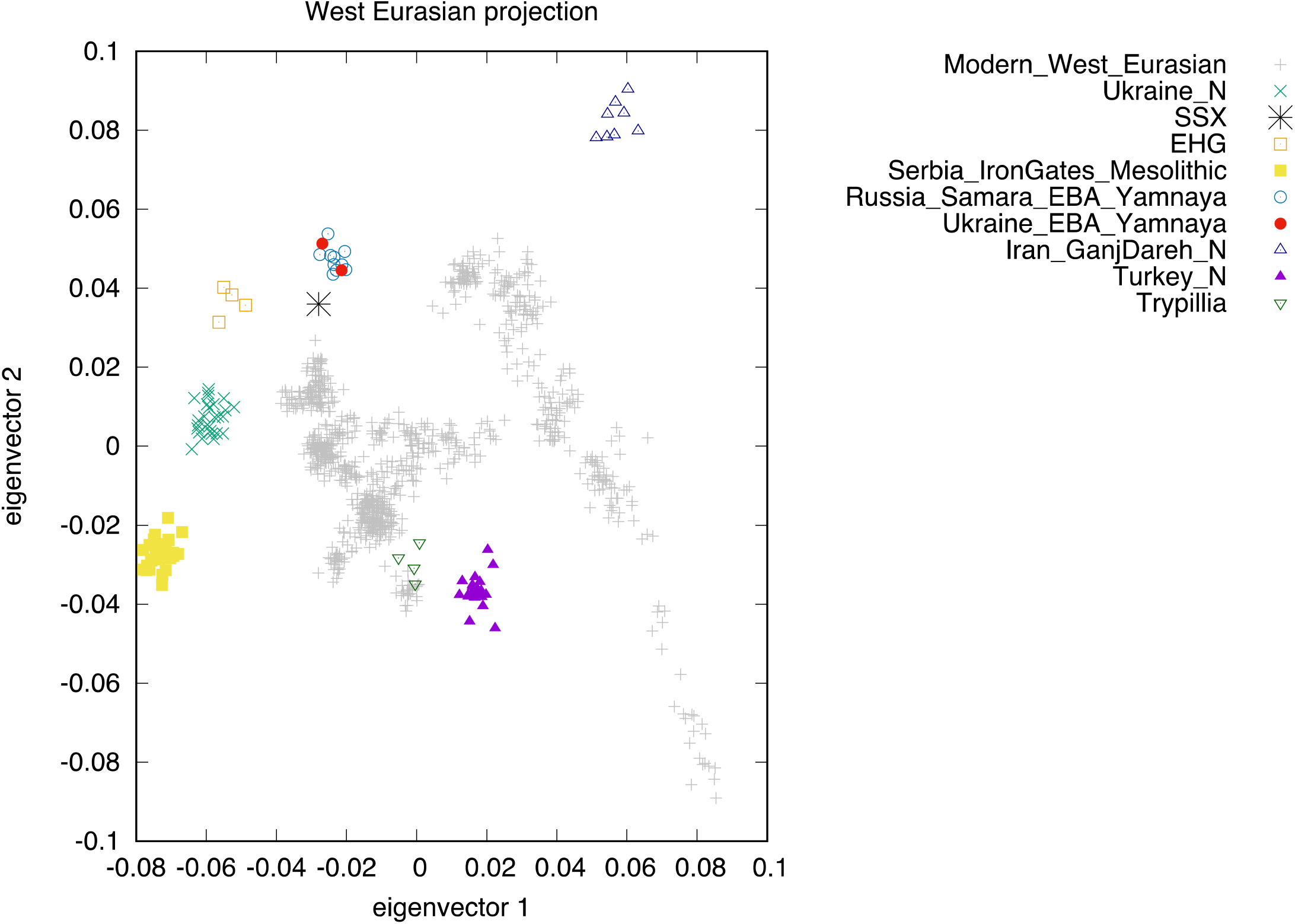
PCA projection of whole genome data of published West Eurasian prehistoric and modern (grey points) populations. Ukraine_N, Ukrainian Neolithic; SSX, KYT specimen; EHG, Eastern Hunter Gatherers; Iran_GanjDareh_N, Iranian Neolithic; Turkey_N, Anatolian Neolithic.

While the KYT-SSX specimen clusters most closely with Yamna, both from Ukraine (Shevchenko) and Russia (Middle Volga, Samara Region), it does not form a clade with the Yamna. An *f_4_* test (Patterson *et al*. 2012) for *f_4_ (Ancient African genomes, Serbia Mesolithic; SSX, Ukraine EBA Yamnaya*) produced a *Z* score of −3.85, and for Samara Yamna instead of KYT-SSX *Z* = −4.78. Statistical analysis of whole-genome data further revealed that KYT-SSX was not genetically similar to Trypillia individuals studied to date. Thus, for example, for *f_4_ (Old Africa, Turkey Neolithic; SSX, Trypillia)* the *Z*-score is 6.692. A plausible scenario is that the KYT individual has much of their ancestry from a Proto-Yamna population (Chintalapati *et al*. 2022), while also being admixed with Serbia/Iron Gates Mesolithic, the latter possibly coming from the Neolithic populations of the Dnipro Valley that have been shown to carry WHG admixture (Mathieson *et al*. 2018). Sredny Stog is the main Proto-Yamna group of the Eneolithic North Pontic steppe and has been hypothesized to be ancestral to Yamna based on archaeological analysis (Telegin 1986), which would be consistent with the assignment of the KYT individual to deriving most of its ancestry from Sredny Stog people.

The presence of an individual with Yamna-related ancestry, most plausibly derived from Sredny Stog groups, at the Trypillian KYT site proposes a possibility of genetic interactions between the steppe and farming worlds as early as the late 5^th^ – early 4^th^ millennium BCE. However, evidence for such interactions is currently lacking. As mentioned above, Trypillians studied to date did not carry steppe genetic admixture until after 3482-3297 calBCE. At the same time, details about Trypillian ancestry of the first part of the 4^th^ millennium come from a single site (Verteba Cave) and a limited number of specimens. Nevertheless, the widespread presence of Cucuteni C pottery at Trypillian sites of the early-middle period, including Verteba, suggests the Sredny Stog-Trypillia interactions were extensive, thus providing multiple opportunities for biological interactions between the two culture groups as well. Further examination of genomes from Sredny Stog and Trypillia sites would help clarify the timing of genetic interactions between Trypillians and Eneolithic steppe forager-pastoralists.

In conclusion, the archaeological, stable isotope and genetic analyses presented in this report produce the following life history highlights for the KYT individual. While the bone fragment of the KYT individual was found in Trypillia culture context, the individual was not genetically associated with the ancestral pool of European Neolithic farmers from which Trypillian ancestry is derived. The KYT individual, genetically a female, is a representative of a Proto-Yamna population of Eneolithic forager-pastoralists of the North Pontic area, such as Sredny Stog. The individual’s maternal (mtDNA) lineage stems from an autochthonous to the North Pontic area clade, which was present in the Dnipro Valley since the beginning of the Holocene. The individual‘s diet was consistent with that of a forager-pastoralist, or the individual had a mixed diet, potentially including resources coming from foraging, pastoralism and farming. The origin of the KYT individual is likely in the middle Dnipro Valley, plausibly from a location in the vicinity of the Deriivka II settlement of Sredny Stog.

Integrating the data presented in this report with the existing body of archaeological knowledge presents the following picture of the population dynamics at the end of the 5^th^ millennium in the East Balkans - North Pontic area. Archaeological evidence indicates the existence of contacts between Sredny Stog, PCCTC and the KGK communities of West Pontic from before 4700-4600 BCE through ca. 4200 BCE, during the operation of the Eneolithic Circum-Pontic trade network (Nikitin & Ivanova 2022). The spread of ceramic finds and copper products allows us to reconstruct the trade route from the western part of KGK culture complex in the Lower Danube Valley (the Pietrele site in Romania, Figure 1) to the Bugeac steppe in the northwest Pontic (e.g. the site of Suvorovo, Figure 1), following north-east to the Middle Dniester PCCTC groups of the Solonceni-Zalischyky type, and then further east to the Dnipro River. Imports of Trypillia BI ceramics at the settlement of Sredny Stog and other archaeological sites in the Lower Dnipro Valley including Mariupol-type cemeteries such as Mykil’s’ke (Nikolskoye) (Figure 1) testify to the extent of these trade connections to the Dnipro Rapids area. This trade route operated for more than 500 years (Videiko 2018). After 4200 BCE, following the collapse of the Balkan Eneolithic in the result of climatic changes (Nikitin & Ivanova 2022) that also affected the steppe belt north of the Black Sea, the Sredny Stog populations moved from the steppe to forest steppe along the Middle Dnipro Valley, where they came into direct contact with PCCTC. This contact is evidenced by the spread of Cucuteni C ceramics, patterned after the steppe ceramics, in Trypillian settlements. It has been suggested that the Cucuteni C ceramic style manifests the presence of the bearers of steppe ceramic technology at Trypillian settlements (Videiko & Burdo 2018). The population affiliation of the KYT individual with Proto-Yamna/Sredny Stog and the presence at KYT of Cucuteni C ceramics made on site not only confirms the presence of Sredny Stog at Trypillian sites, but also reflects the integration of steppe individuals into Trypillian societies. This study exemplifies a case when the hypotheses of archaeologists, based on the use of traditional typological and comparative methods, received strong validation from molecular analysis methods employed by natural sciences.

## Acknowledgements

The authors would like to thank Nadin Rohland, Swapan Mallick, Matthew Mah, Adam Micco, and Aisling Kearns for their skillful preparation and processing of the sample for ancient DNA analysis. This study was supported, in part, by Professional Development Funds from the College of Liberal Arts and Sciences of Grand Valley State University to AGN and by the Howard Hughes Medical Institute (HHMI) to DR. The acquisition of the Nu Plasma II MC-ICP-MS was funded by the National Science Foundation (grant #BCS-0922374). The Archaeometry Laboratory is supported by the National Science Foundation (grant #BCS-1912776). This article is subject to the HHMI’s Open Access to Publications policy. HHMI lab heads have previously granted a nonexclusive CC BY 4.0 license to the public and a sublicensable license to HHMI in their research articles. Pursuant to those licenses, the author-accepted manuscript of this article can be made freely available under a CC BY 4.0 license immediately upon publication.

