## Supplemental Figures S1-5 for "Interactions between Trypillian farmers and North Pontic forager-pastoralists in Eneolithic central Ukraine"

Figures S1-S5

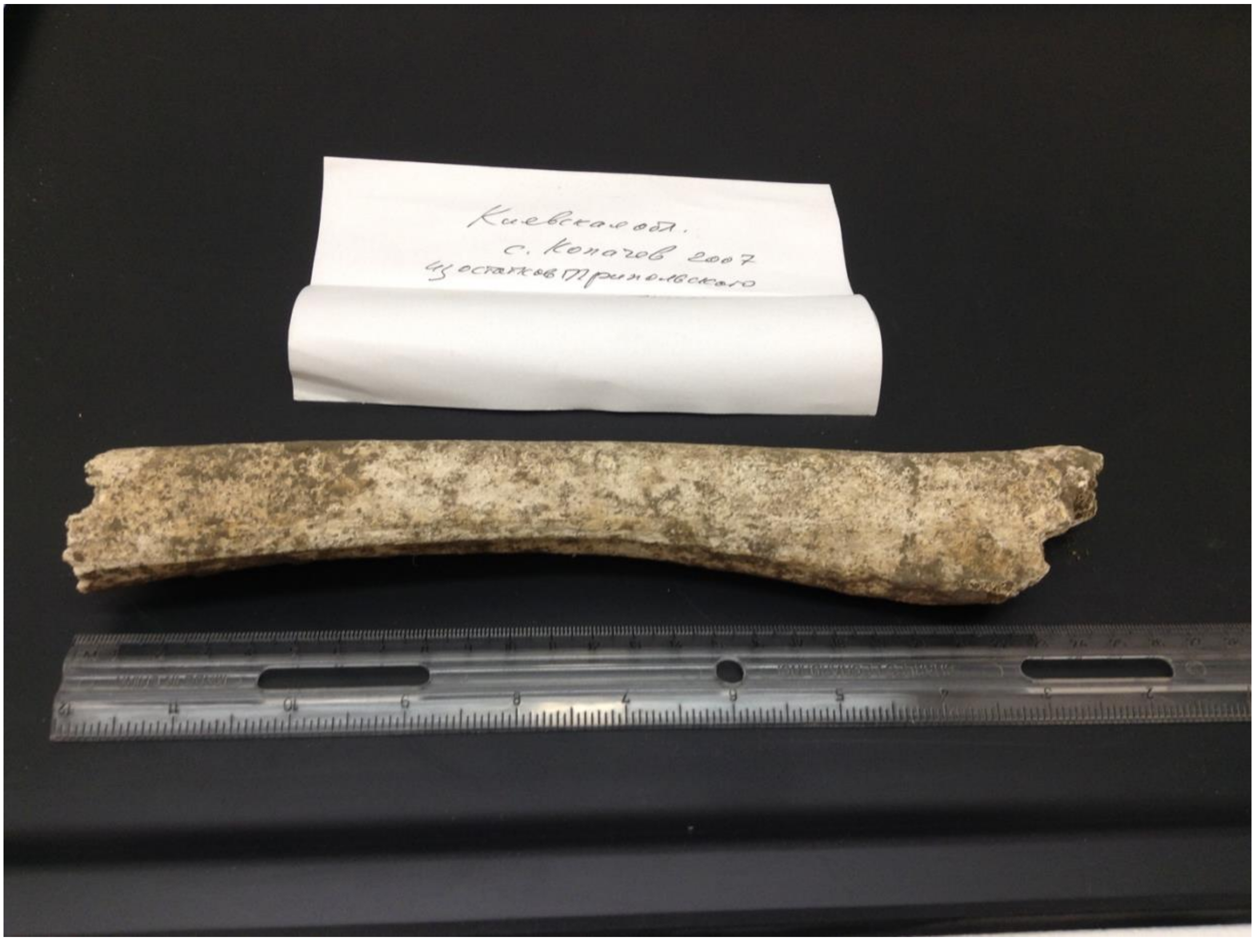


**Figure S1.** Long bone fragment from the Trypillian layer at the Kolomiytsiv Yar Tract. Photo A. G. Nikitin, 2016

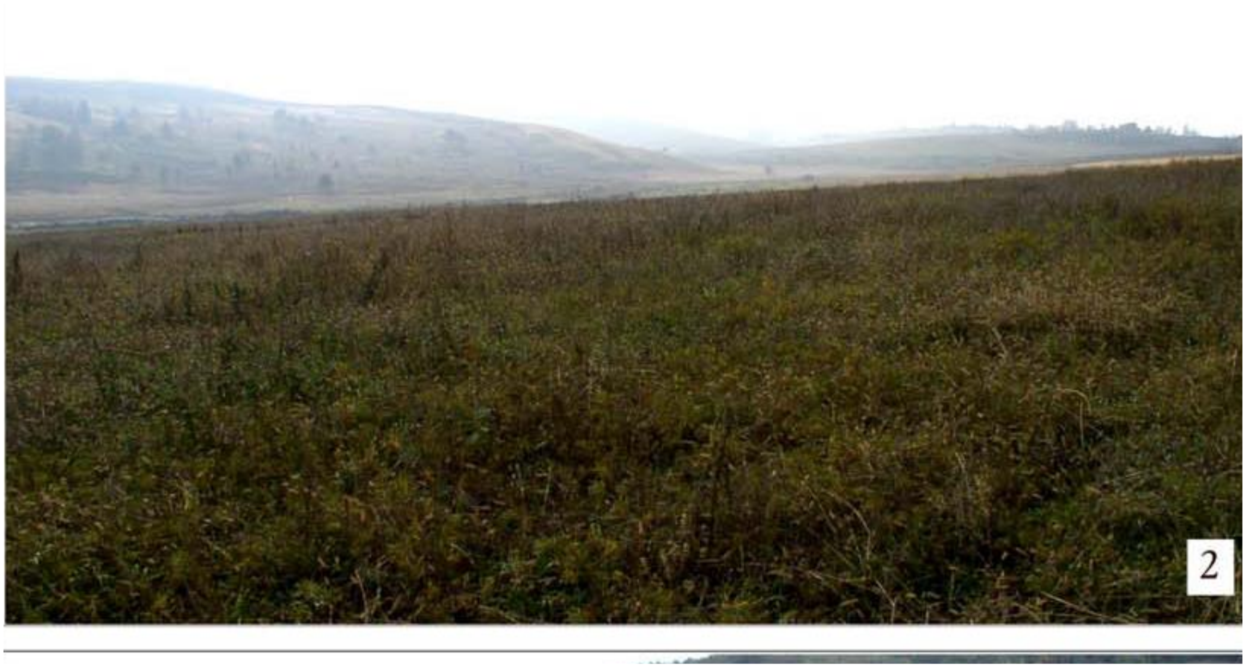


**Figure S2**. The Kolomiytsiv Yar Tract location. Photo by M. Y. Videiko, 2006.

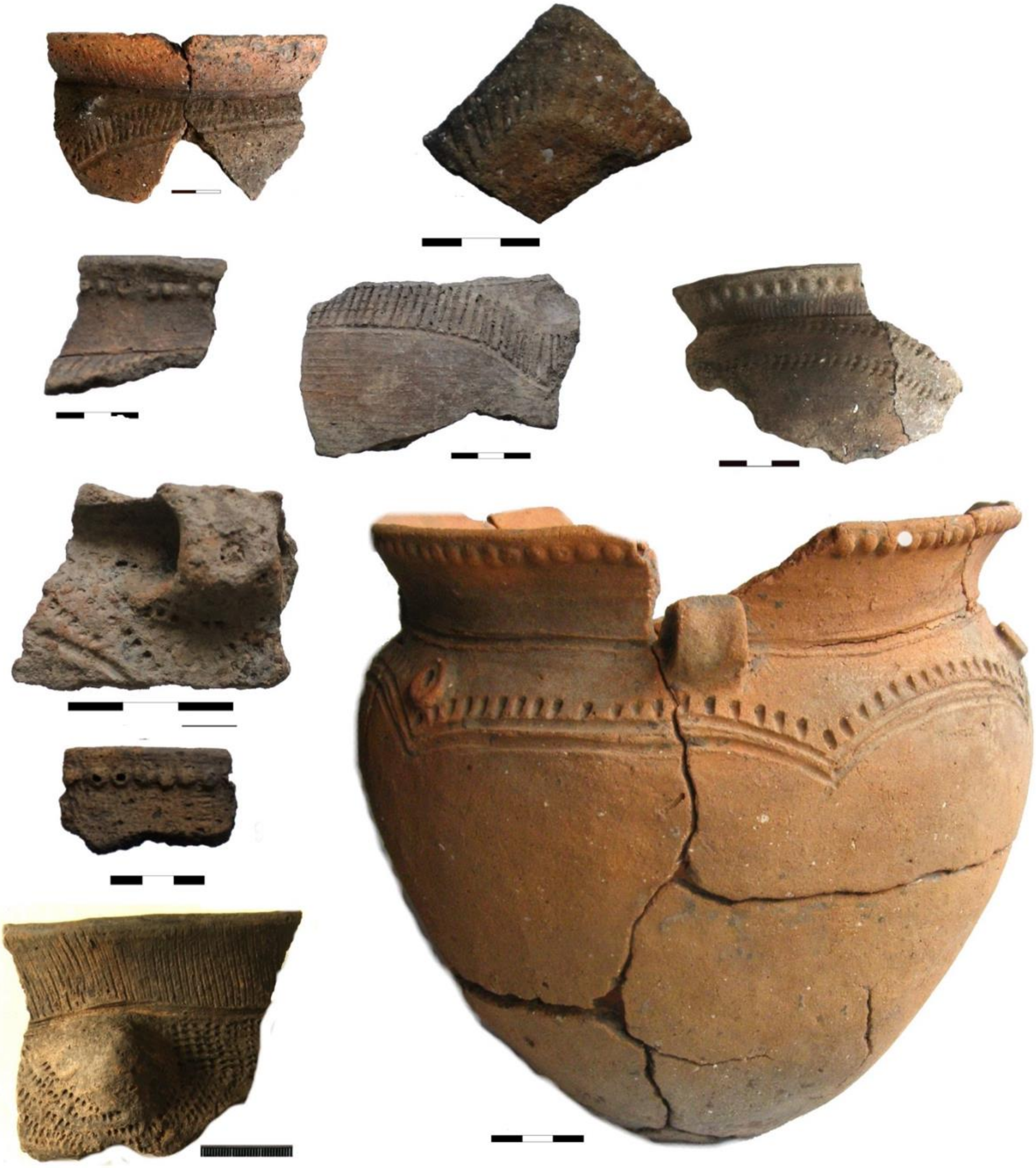

**Figure S3**. Cucuteni C pottery from the Trypillian settlement of Kolomiytsiv Yar Tract.

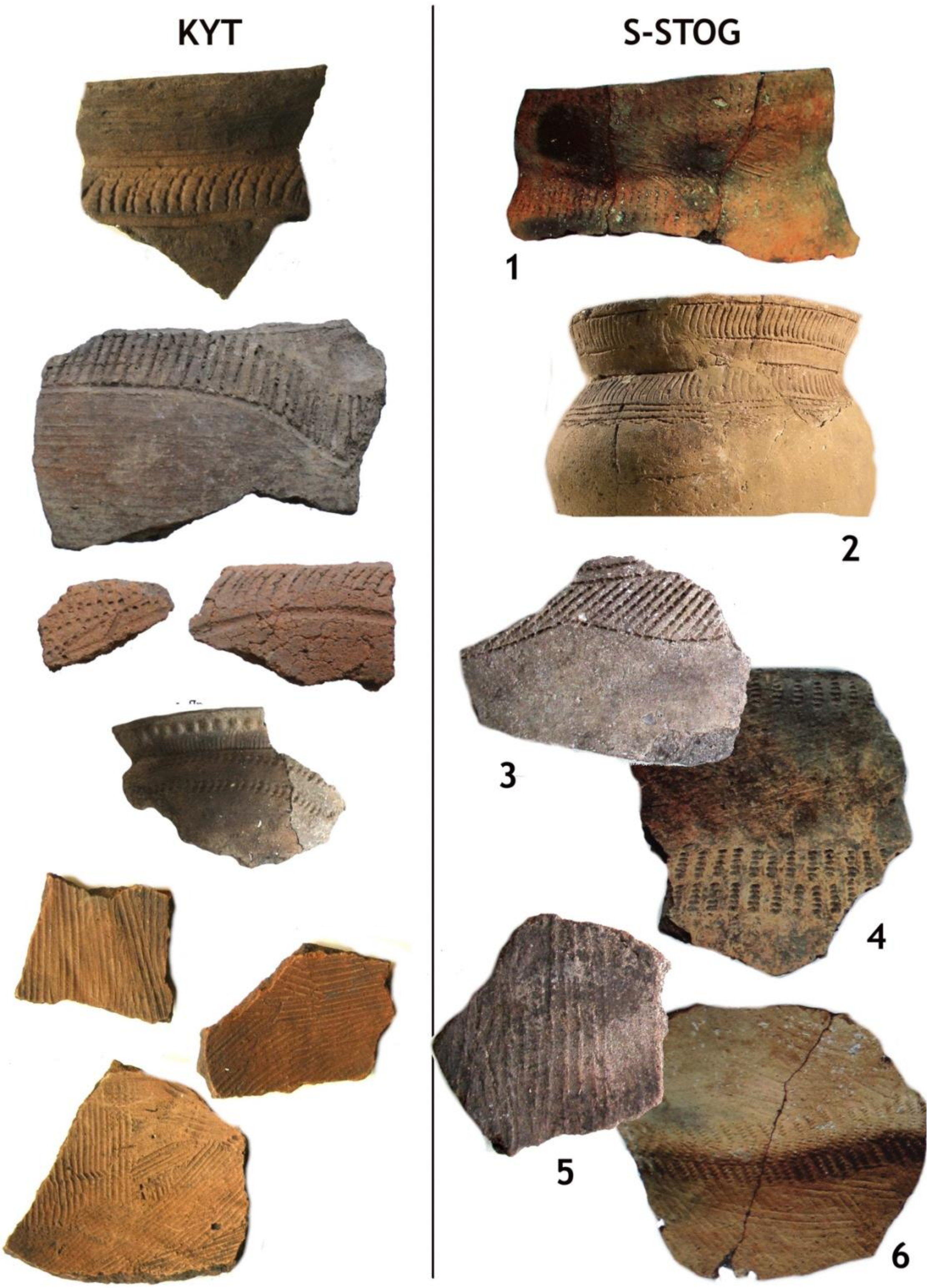


**Figure S4**. Cucuteni C pottery from Kolomiytsiv Yar Tract (KYT) and Sredny Stog (S-STOG) culture sites. 1,4,6 - Molyukhiv Bugor (after T. Neradenko); 2- Sredny Stog II; 3,5- Stril'cha Skelya.

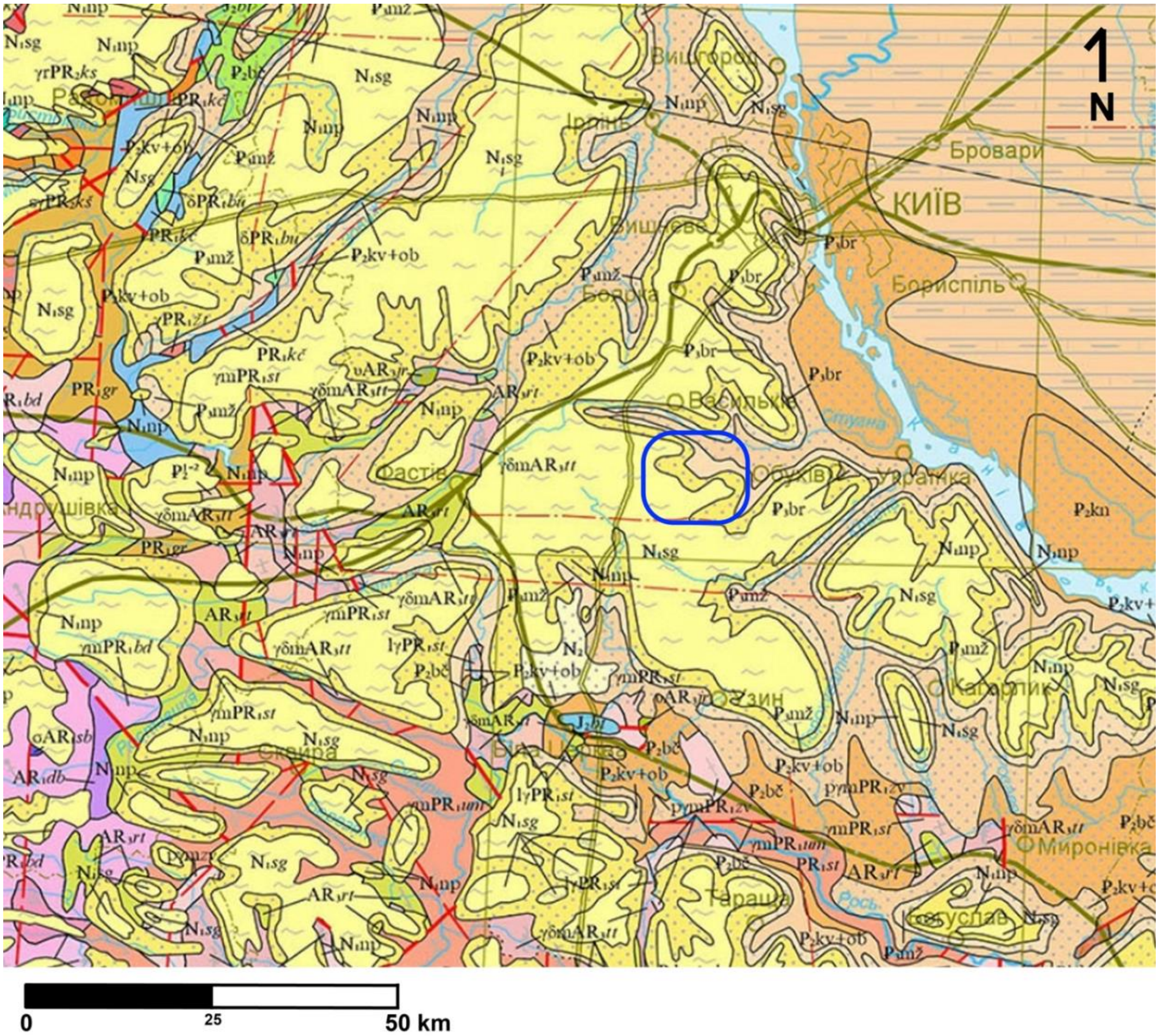


**Figure S5**. Regional lithology in the vicinity of the Kolomiytsiv Yar Tract. Image from https://minerals-ua.info/w/mapviewe.php?pr=2 (modified).

The site (outlined in blue) is situated along a small drainage that likely cuts through the following geological units/lithologies:

- $N_1$np (Miocene) Novopetrivsky Regional Stage – sands, sandstones, clays, including bentonitic and fireproof clays, bands of brown coal
- $P_3$mz (Oligocene) Mezhyhirsky Regional Stage – glauconitic quartz, argillaceous sands, sometimes – with phosphorite strips
- $P_2$kv+ob (Eocene) Kyiv and Obukiv Regional Stages – marls, clays, glauconite sands, aleurites, sandstones, including opoka-like sandstones

See https://minerals-ua.info/w/mapviewe.php?pr=2 for detailed legend.
